# Evolution of pre-copulatory and post-copulatory strategies of inbreeding avoidance and associated polyandry

**DOI:** 10.1101/136093

**Authors:** A. Bradley Duthie, Greta Bocedi, Ryan R. Germain, Jane M. Reid

## Abstract

Inbreeding depression is widely hypothesised to drive adaptive evolution of pre-copulatory and post-copulatory mechanisms of inbreeding avoidance, which in turn are hypothesised to affect evolution of polyandry (i.e., female multiple mating). However, surprisingly little theory or modelling critically examines selection for pre-copulatory or post-copulatory inbreeding avoidance, or both strategies, given evolutionary constraints and direct costs, or examines how evolution of inbreeding avoidance strategies might feed back to affect evolution of polyandry. Selection for post-copulatory inbreeding avoidance, but not for pre-copulatory inbreeding avoidance, requires polyandry, while interactions between pre-copulatory and post-copulatory inbreeding avoidance might cause functional redundancy (i.e., ‘degeneracy’) potentially generating complex evolutionary dynamics among inbreeding strategies and polyandry. We used individual-based modelling to quantify evolution of interacting pre-copulatory and post-copulatory inbreeding avoidance and associated polyandry given strong inbreeding depression and different evolutionary constraints and direct costs. We found that evolution of post-copulatory inbreeding avoidance increased selection for initially rare polyandry, and that evolution of a costly inbreeding avoidance strategy became negligible over time given a lower cost alternative strategy. Further, fixed pre-copulatory inbreeding avoidance often completely precluded evolution of polyandry and hence post-copulatory inbreeding avoidance, but fixed post-copulatory inbreeding avoidance did not preclude evolution of pre-copulatory inbreeding avoidance. Evolution of inbreeding avoidance phenotypes and associated polyandry are therefore affected by evolutionary feedbacks and degeneracy. All else being equal, evolution of pre-copulatory inbreeding avoidance and resulting low polyandry is more likely when post-copulatory inbreeding avoidance is precluded or costly, and evolution of post-copulatory inbreeding avoidance greatly facilitates evolution of costly polyandry.

## Introduction

Inbreeding, defined as reproduction between relatives, often greatly reduces the fitness of resulting inbred offspring (termed ‘inbreeding depression’; Charlesworth & Charlesworth, 1999; Keller & Waller, 2002; Charlesworth & Willis, 2009). Such strong inbreeding depression is widely hypothesised to drive evolution of inbreeding avoidance, which can be enacted through multiple reproductive strategies (Parker, 1979, 2006; Pusey & Wolf, 1996; Szulkin *et al.*, 2013).

From a female’s perspective, inbreeding avoidance might be achieved by avoiding mating with related males (i.e., pre-copulatory inbreeding avoidance), or by biasing fertilisation towards unrelated males following mating (i.e., post-copulatory inbreeding avoidance). Evolution of post-copulatory inbreeding avoidance requires that females express some degree of polyandry, defined as mating with multiple males during a single reproductive bout (but see Dougherty *et al.*, 2016). Further, such polyandry might itself evolve because it allows females to mate with additional unrelated males following an initial mating with a relative, potentially including males that were unavailable for initial mate choice (e.g., Reid *et al.*, 2015*b*; Duthie *et al.*, 2016). Polyandry can thereby facilitate pre-copulatory inbreeding avoidance even without any post-copulatory female choice or otherwise biased fertilisation among sperm (i.e., under conditions of a ‘fair raffle’).

Overall, polyandry can therefore simultaneously allow females to mate with less closely related males and create opportunity for further inbreeding avoidance enacted through active post-copulatory choice. Indirect selection on polyandry resulting from reduced inbreeding depression in offspring fitness could help explain evolution of polyandry in cases where multiple mating decreases female reproductive success, imposing a direct cost on polyandrous females (Zeh & Zeh, 1997; Jennions & Petrie, 2000; Tregenza & Wedell, 2002). However, despite such widely-invoked hypotheses, there is surprisingly little theory or modelling that critically examines the conditions under which pre-copulatory or post-copulatory inbreeding avoidance, or both strategies, are predicted to evolve, or that examines how evolution of such strategies might feed back to affect underlying evolution of polyandry. Comprehensive understanding of evolution of reproductive strategies given inbreeding depression requires consideration of the fundamental joint effects of selection on pre-copulatory and post-copulatory inbreeding avoidance and polyandry.

Despite the paucity of theory, numerous empirical studies on diverse species have tested for, and in some cases found evidence of, female inbreeding avoidance in systems where polyandry also occurs (Tregenza & Wedell, 2002; Varian-Ramos & Webster, 2012; Kingma *et al.*, 2013; Arct *et al.*, 2015, but see Reid 2015). However, few studies have determined whether inbreeding avoidance is enacted through pre-copulatory or post-copulatory mechanisms. Among these studies, pre-copulatory inbreeding avoidance has been reported in sweet potato weevils (*Cylas formicarius*; Kuriwada *et al.*, 2011), purple-crowned fairy-wrens (*Malurus coronatus*; Kuriwada *et al.*, 2011), and squinting bush brown butterflies (*Bicyclus anynana*; Fischer *et al.*, 2015), while post-copulatory inbreeding avoidance has been reported in, for example, red junglefowl (*Gallus gallus*; Pizzari *et al.*, 2004) and crickets (*Teleogryllus oceanicus*, *Gryllus bimaculatus*; Simmons *et al.*, 2006; Bretman *et al.*, 2009). Evidence for both pre-copulatory and post-copulatory inbreeding avoidance is available across different studies of Trinidadian guppies (*Poecilia reticulata*; Gasparini & Pilastro, 2011; Daniel & Rodd, 2015) and house mice (*Mus domesticus*; Potts *et al.*, 1991; Firman & Simmons, 2015). Meanwhile, Liu *et al.* (2014) found evidence of pre-copulatory but not post-copulatory inbreeding avoidance within a single study of cabbage beetles (*Colaphellus bowringi*). However, Ala-Honkola *et al.* (2011) and Tan *et al.* (2012) found no evidence for pre-copulatory or post-copulatory inbreeding avoidance in fruit flies (*Drosophila melanogaster*), respectively, and Reid et al. (2015*a*; 2015*b*) found no net inbreeding avoidance in song sparrows (*Melospiza melodia*) despite strong inbreeding depression and opportunities for both pre-copulatory and post-copulatory inbreeding avoidance. Taken together, these studies demonstrate that diverse combinations of pre-copulatory and post-copulatory inbreeding avoidance, or lack thereof, occur in nature. However, there is as yet no theory that predicts what combinations of pre-copulatory and post-copulatory inbreeding avoidance and associated polyandry should be favoured by selection when all can evolve. Consequently, there is no theory that allows the diversity of observed pre-copulatory and post-copulatory inbreeding avoidance strategies to be interpreted, and there are no clear hypotheses that could be tested through future empirical studies of individual systems or subsequent comparative analyses.

In one first step, Duthie *et al.* (2016) used a genetically-explicit individual-based model to examine conditions under which polyandry is predicted to evolve due to selection stemming from pre-copulatory inbreeding avoidance in the absence of post-copulatory inbreeding avoidance. Simulations showed that even when selection for pre-copulatory inbreeding avoidance was strong and females consequently preferred unrelated mates, selection for polyandry specifically to facilitate this inbreeding avoidance occurred only under highly restricted conditions. Key requirements were that direct negative selection (i.e., ‘costs’) on polyandry was weak, that very few males were available for a female’s initial mate choice but many were available for additional mate choice(s), or that polyandry was conditionally expressed when a focal female was related to her initial mate (Duthie *et al.*, 2016). Without these conditions, polyandrous females tended to increase rather than decrease their overall degree of inbreeding, ultimately reducing offspring fitness. This increase occurred because, once pre-copulatory inbreeding avoidance evolved, polyandrous females had already chosen available unrelated males as their initial mates. Their additional mates, chosen from the remaining male population, were therefore increasingly likely to include relatives. Evolution of polyandry purely to facilitate pre-copulatory inbreeding avoidance was consequently restricted (Duthie *et al.*, 2016).

However, if post-copulatory inbreeding avoidance could evolve alongside pre-copulatory inbreeding avoidance, then polyandrous females could bias fertilisation towards unrelated males within their set of mates. Evolution of post-copulatory inbreeding avoidance might consequently reduce the cost of polyandry stemming from the accumulation of related mates across multiple matings, potentially facilitating evolution of polyandry to avoid inbreeding under broader conditions, and driving further evolution of pre-copulatory or post-copulatory mate choice strategies. Yet, if polyandry and pre-copulatory and post-copulatory inbreeding avoidance can all evolve, the long-term dynamics of these three reproductive strategies become difficult to predict. Strong inbreeding depression might drive initial evolution of both pre-copulatory and post-copulatory inbreeding avoidance and associated polyandry. However, the co-occurrence of pre-copulatory and post-copulatory inbreeding avoidance might cause some degree of ‘degeneracy’, defined as a phenomenon by which different elements of a system result in identical outputs (Edelman & Gally, 2001). Consequently, if evolution of polyandry and post-copulatory inbreeding avoidance renders pre-copulatory inbreeding avoidance functionally redundant, or vice versa, then only one inbreeding avoidance strategy might be maintained in the long-term.

The few previous models that considered evolution of biparental inbreeding avoidance through mate choice (as opposed to dispersal) have implicitly or explicitly considered the fate of a rare allele underlying pre-copulatory inbreeding avoidance in a population initially fixed for random mating (e.g., Parker, 1979, 2006; Duthie *et al.*, 2016; Duthie & Reid, 2016). Such models are useful for isolating the invasion fitness of this single strategy. However, when both pre-copulatory and post-copulatory strategies can affect the realised degree of inbreeding, it cannot be assumed that both strategies will simultaneously invade a randomly mating population, nor that the invasion fitness of one strategy will be independent of the pre-existence or invasion fitness of the other strategy. For example, if pre-adaptation or a selective sweep results in fixation of alleles underlying pre-copulatory inbreeding avoidance, then new alleles underlying polyandry and post-copulatory inbreeding avoidance might be unlikely to invade a population because the phenotypic effect of such alleles on the overall degree of inbreeding, and resulting indirect selection, could be negligible. Conversely, fixation of alleles underlying polyandry and post-copulatory inbreeding avoidance might reduce or eliminate positive selection on alleles underlying pre-copulatory inbreeding avoidance and hence impede adaptive evolution of mate choice. New theory, guided by modelling that evaluates invasion dynamics of alleles underlying multiple interacting and potentially functionally redundant (i.e., ‘degenerate’) traits, is therefore needed.

In the context of inbreeding depression as a key hypothesised driver of reproductive strategy evolution, the absolute and relative frequencies of alleles underlying pre-copulatory and post-copulatory inbreeding avoidance and polyandry will be affected not only by the magnitudes of positive indirect selection stemming from reduced inbreeding depression in females’ offspring, but also by the magnitudes of direct negative selection on resulting phenotypes (i.e., the direct fitness costs of expressing each reproductive strategy). Empirical studies have demonstrated diverse costs of mating and mate choice, for example, including energetic costs of developing, maintaining, or enacting necessary physiologies (e.g., Gasparini & Pilastro, 2011; Tuni *et al.*, 2013; Fitzpatrick & Evans, 2014); increased risks of predation or disease stemming from increased mate-searching or mating (e.g., Rowe, 1988; Ronkainen & Ylonen, 1994; Koga *et al.*, 1998); increased risk of complete mating or fertilisation failure given extreme choosiness (Kokko & Mappes, 2013); and risks of harm stemming from sexual conflict over fertilisation (e.g., Rowe *et al.*, 1994). If the relative costs of pre-copulatory and post-copulatory inbreeding avoidance differ, then alleles underlying the less costly strategy might become fixed over generations, while alleles underlying the more costly strategy might go extinct, especially if their effects become redundant following evolution of the less costly strategy. Dynamic models that track the frequencies of alleles underlying multiple, potentially interacting, inbreeding avoidance strategies that are enacted among relatives resulting from reproductive strategies and inbreeding depression expressed in previous generations are consequently useful to understand and predict evolutionary outcomes.

We use individual-based modelling to address three key questions regarding evolution of pre-copulatory and post-copulatory inbreeding avoidance and associated polyandry given opportunity for inbreeding and strong inbreeding depression. First, does evolution of post-copulatory inbreeding avoidance, alongside pre-copulatory inbreeding avoidance, facilitate evolution of polyandry? Second, how do costs associated with polyandry and pre-copulatory and post-copulatory inbreeding avoidance affect evolutionary outcomes and, in particular, the long-term persistence of these reproductive strategies given cost asymmetry? Third, how is selection for initially rare pre-copulatory or post-copulatory inbreeding avoidance affected if the other strategy of inbreeding avoidance is already fixed? To address these questions, we designed our model to isolate the effect of each biological mechanism of interest (i.e., post-copulatory inbreeding avoidance, cost asymmetry, and strategy pre-existence, respectively) and hence to allow comparison of simulations with the mechanism present versus absent with otherwise identical parameter values and conditions. We thereby illustrate how the simultaneous evolution of multiple interacting degenerate reproductive strategies can generate diverse evolutionary outcomes.

## Model

We model evolution of polyandry, and of pre-copulatory and post-copulatory inbreeding avoidance strategies (hereafter simply ‘inbreeding strategies’ because the model did not preclude evolution of inbreeding preference), in a small focal population by tracking the dynamics of alleles underlying reproductive strategies expressed by females. We thereby track evolutionary dynamics given internally consistent patterns of relatedness caused by non-random mating and capture effects of mutation, gene flow, drift, and selection.

Complex traits such as reproductive strategies are likely to be polygenic (Evans & Simmons, 2008). Hence, we model individuals with 10 physically unlinked diploid loci (i.e., 20 alleles), underlying each of three reproductive strategy traits: tendency for polyandry (*P*_*a*_, ‘polyandry’ alleles), pre-copulatory inbreeding strategy (*M*_*a*_, ‘mating’ alleles), and post-copulatory inbreeding strategy (*F*_*a*_, ‘fertilisation’ alleles). All individuals therefore have 30 diploid loci (i.e., 60 alleles) in total, each of which can take the value of any real number (continuum-of-alleles model; Kimura, 1965; Lande, 1976; Reeve, 2000; Bocedi & Reid, 2014).

Alleles combine additively to determine genotypic values (*P*_*g*_, *M*_*g*_, and *F*_*g*_) and resulting phenotypic values (*P*_*p*_, *M*_*p*_, and *F*_*p*_) for tendency for polyandry, pre-copulatory inbreeding strategy, and post-copulatory inbreeding strategy, respectively. Each individual’s genotypic values *P*_*g*_, *M*_*g*_, and *F*_*g*_ equal the sum of its 20 alleles for each trait. Each individual’s phenotypic values for pre-copulatory and post-copulatory inbreeding strategy equal their respective genotypic values (*M*_*p*_ = *M*_*g*_ and *F*_*p*_ = *F*_*g*_), where negative and positive values cause inbreeding avoidance and preference, respectively (see details of mating and fertilisation strategies below).

In contrast, individuals’ phenotypic values for tendency for polyandry (*P*_*p*_) cannot map directly onto their genotypic values (*P*_*g*_) because *P*_*g*_ can evolve to be negative, but females cannot mate with a negative number of additional males (e.g., Shuker *et al.*, 2007; Evan & Gasparini, 2013). Rather, we considered polyandry “threshold trait”, whereby continuous genotypic variation translates into expression of discrete phenotypic value(s) at some threshold (Lynch & Walsh, 1998; Roff, 1996, 1998; Duthie *et al.*, 2016). Accordingly, we allow individuals’ phenotypic values for tendency for polyandry to equal genotypic values (*P*_*p*_ = *P*_*g*_) if *P*_*g*_ ≥ 0, but set *P*_*p*_ = 0 if *P*_*g*_ < 0. A negative *P*_*g*_ value therefore generates a female that is phenotypically monandrous, while a positive *P*_*g*_ value generates a female that can express some degree of polyandry (see details below). Polyandry is therefore influenced by continuous genetic variation but only expressed when *P*_*g*_ > 0. One important general property of such threshold traits is that deleterious traits are occasionally expressed despite sustained negative selection because recombination among deleterious alleles can cause the underlying genotypic value to exceed the threshold for expression (Roff, 1996, 1998).

In overview, each model generation proceeds with females paying costs associated with their reproductive strategy traits, and expressing polyandry, mating, and fertilisation. Offspring inherit a randomly sampled allele from each parent at each locus. Alleles can then mutate and offspring express inbreeding depression in viability. Immigrants arrive in the population and density regulation limits population growth (see below). We record the population pedigree and directly calculate the coefficient of kinship (*k*) between all potential mates in each generation (defined as the probability that two homologous alleles will be identical-by-descent, therefore ranging from 0-1), allowing individual pre-copulatory and post-copulatory inbreeding strategies to be enacted. Values of *k* are calculated directly from the pedigree using a standard iterative algorithm (e.g., ?Duthie et al., 2016). Key individual traits, parameter values, and variables are described in Table 1.

**Table 1:** Individual traits (A), model parameter values (B), and model variables (C) for an individual-based model of the evolution of polyandry, pre-copulatory inbreeding strategy, and post-copulatory inbreeding strategy.

| A | Trait | Allele | Genotype | Phenotype |
| --- | --- | --- | --- | --- |
| | Tendency for polyandry | $P_a$ | $P_g$ | $P_p$ |
| | Pre-copulatory inbreeding strategy | $M_a$ | $M_g$ | $M_p$ |
| | Post-copulatory inbreeding strategy | $F_a$ | $F_g$ | $F_p$ |

| B | Description | Parameter | Default value(s) |
| --- | --- | --- | --- |
| | Cost of tendency for polyandry | $c_P$ | 0, 0.02 |
| | Cost of pre-copulatory inbreeding strategy | $c_M$ | 0, 0.02 |
| | Cost of post-copulatory inbreeding strategy | $c_F$ | 0, 0.02 |
| | Focal female's number of offspring | $n$ | 8 |
| | Log-linear slope of inbreeding depression | $\beta$ | 3 |
| | Adult male immigrants per generation | $\rho$ | 5 |
| | Female carrying capacity | $K_f$ | 100 |
| | Male carrying capacity | $K_m$ | 100 |
| | Mutation rate of alleles | $\mu$ | 0.001 |
| | Standard deviation of mutation effect size | $\mu_{SD}$ | $\sqrt{1/20}$ |

| C | Description | Variable |
| --- | --- | --- |
| | Coefficient of kinship | $k$ |
| | Focal female's number of mates | $N_{males}$ |
| | Female $i$'s perceived absolute mate quality of male $j$ | $Q_{ij}^m$ |
| | Female $i$'s perceived relative mate quality of male $j$ | $q_{ij}^m$ |
| | Female $i$'s perceived absolute fertilisation quality of male $j$ | $Q_{ij}^f$ |
| | Female $i$'s perceived relative fertilisation quality of male $j$ | $q_{ij}^f$ |
| | Viability of a focal female's offspring | $\Psi_{off}$ |

### Costs

Phenotypic values of the three reproductive strategy traits each incur set costs that combine to independently increase the probability that a focal female will die before mating (realisation of costs precedes mating and fertilisation, so we present further details of mating and fertilisation below). Numerous forms and mechanisms of direct costs of reproductive strategies could be hypothesised and modelled; the most appropriate formulation depends on the question (see Discussion). By allowing costs of the three strategies to be directly and independently controlled, our model facilitates direct comparison of evolution of pre-copulatory versus post-copulatory inbreeding avoidance given known cost asymmetries. Qualitatively, such costs on female survival probability are biologically reasonable. For example, polyandrous females that undertake increased mate searching or mating can experience increased predation risk (e.g., Rowe, 1988; Ronkainen & Ylonen, 1994; Koga *et al.*, 1998). Females that express pre-copulatory choice can increase the risk of mortality due to harm caused by sexual conflict over mating, and also risk complete mating failure (which equates to pre-reproductive mortality in semelparous organisms; Rowe *et al.*, 1994; Kokko & Mappes, 2013). Finally, females that express post-copulatory choice can pay up-front energetic costs, which might result in trade-offs with survival due to developing physiological or biochemical mechanisms needed to store sperm and successfully bias fertilisation (e.g., Gasparini & Pilastro, 2011; Tuni *et al.*, 2013; Fitzpatrick *et al.*, 2014).

Accordingly, the probabilities of pre-mating mortality due to the costs of polyandry (*c*_*P*_), pre-copulatory inbreeding strategy (*c*_*M*_), and post-copulatory inbreeding strategy (*c*_*F*_) are *P*_*p*_ × *c*_*P*_, |*M*_*p*_| × *c*_*M*_, and |*F*_*p*_| × *c*_*F*_, respectively. Here |*M*_*p*_| and |*F*_*p*_| are the absolute values of *M*_*p*_ and *F*_*p*_, respectively. Absolute values are used for applying costs to inbreeding avoidance strategies because both negative and positive *M*_*p*_ and *F*_*p*_ values could potentially arise and affect the degree of inbreeding, representing inbreeding avoidance and inbreeding preference, respectively. In contrast, *P*_*p*_ is already defined to be non-negative. Overall, because generations are non-overlapping, a female’s probability of total reproductive failure increases linearly with the phenotypic value of each trait.

### Mating and pre-copulatory inbreeding avoidance

After costs are realised, each surviving female chooses *N*_*males*_ males to mate with, where *N*_*males*_ is calculated by sampling from a Poisson distribution such that *N*_*males*_ = *Poisson*(*P*_*p*_) + 1. This ensures that all surviving females choose at least one mate and generates each female’s realised degree of polyandry with some stochastic variation around the expected mean *N*_*males*_ of *P*_*p*_ + 1.

All males in the population are assumed to be available for any female to choose. We therefore assume that there is no opportunity cost of male mating, so mating with one female does not reduce a male’s availability to mate with any other female. Females mate with *N*_*males*_ without replacement, meaning that *N*_*males*_ models a female’s total number of different mates rather than solely her total number of matings.

Most often, *N*_*males*_ will be smaller than the total number of available males (Duthie *et al.*, 2016). Each female then chooses her *N*_*males*_ mates based on her pre-copulatory inbreeding avoidance phenotype (*M*_*p*_). Negative or positive *M*_*p*_ values cause a female to avoid or prefer mating with kin, respectively, whereas *M*_*p*_ = 0 causes a female to mate randomly with respect to kinship.

To calculate the probability that a female *i* mates with a male *j* to whom she is related by some kinship *k*_*ij*_, each male is first assigned a perceived mate quality 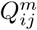. If the female has a strategy of pre-copulatory inbreeding avoidance (*M*_*p*_ < 0), then 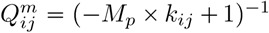, meaning that 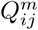 decreases linearly with increasingly positive alues of *k*_*ij*_ and increasingly negative values of *M*_*p*_. If the female has a strategy of pre-copulatory inbreeding preference (*M*_*p*_ > 0), then 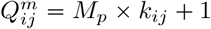, meaning that 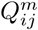 increases with increasingly positive *k*_*ij*_ and *M*_*p*_. If *M*_*p*_ = 0, then all males are assigned 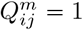.

Each male’s value with respect to a female *i* is then divided by the sum of all 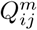 values across all males with respect to that female, thereby assigning each male a relative perceived quality 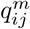, which is constrained to values between zero and one. The value of 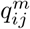 then defines the probability that female *i* mates with male *j*. Mating is therefore stochastic, and females do not always mate with the male of the highest 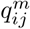. For polyandrous females that choose multiple mates (i.e., *N*_*mates*_ > 1), mates are chosen iteratively such that 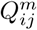 and 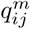 are re-calculated for each additional mate choice, and with 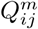 and therefore 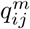 values of already chosen males set to zero to ensure mate sampling without replacement. In the unlikely event that a female’s *N*_*males*_ exceeds the total number of available males, then she simply mates with all males.

### Fertilisation and post-copulatory inbreeding avoidance

Following mating, fertilisation occurs such that each of a female’s *n* offspring is independently assigned a sire (with replacement) from the *N*_*males*_ with which the female mated. Sire identity depends on female’s kinship with each mate (*k*_*ij*_) and her post-copulatory inbreeding strategy phenotype (*F*_*p*_). Negative and positive values of *F*_*p*_ correspond to post-copulatory inbreeding avoidance or preference, respectively, whereas *F*_*p*_ = 0 causes random fertilisation with respect to kinship.

The probability that an offspring of female *i* is sired by any one of her mates *j* is calculated by assigning a perceived fertilisation quality to each *j* with respect to 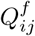. Perceived fertilisation quality 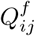 is calculated in the same way as perceived mate quality 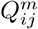, such that if female *i* has a strategy of post-copulatory inbreeding avoidance (*F*_*p*_ < 0), then the perceived quality of male *j* is 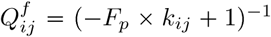. If the female has a strategy of post-copulatory inbreeding preference (*F*_*p*_ > 0), then the perceived quality of male *j* is 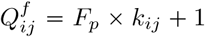. A relative quality 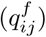 is then calculated for each male by dividing his 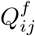 by the sum of the 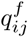 values across all of a female’s mates. These 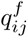 values, which lie between zero and one, define the probability of paternity. Females produce *n* offspring, so a female *i* samples from her mates *n* times independently and with replacement with a probability of for each male *j* to determine the realised distribution of sires. Offspring have equal probability of being female or male. After offspring production, all female and male adults die so that generations are non-overlapping.

### Mutation

Offsprings’ alleles mutate with independent probabilities *μ* = 0.001. When a mutation occurs, a mutation effect size is sampled from a normal distribution with a mean of zero and a standard deviation of *μ*_*SD*_ and added to the original allele value (Kimura, 19<u>65; L</u>ande, 1976; Bocedi & Reid, 2014; Duthie *et al.*, 2016). The value of *μ*_*SD*_ is set to 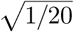.

### Inbreeding depression

The viability of a female *i*’s offspring (Ψ_off_) decreases as a log-linear function of her kinship with the sire *j* of the offspring (*k*_*ij*_) and inbreeding depression slope *β*,

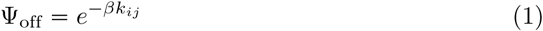

Here, *β* models the number of haploid lethal equivalents that exist as deleterious recessive alleles in the gametes of *i* and *j*, and which might be homozygous in offspring and reduce viability. Equation 1 assumes independent allelic effects, generating multiplicative effects on offspring viability (Morton *et al.*, 1956; Mills & Smouse, 1994). It also assumes that inbreeding does not covary with inbreeding load (i.e., no purging). This formulation ensures that the relationship between *k*_*ij*_ and the magnitude of inbreeding depression in offspring is consistent across replicate simulations. This choice, as opposed to a more mechanistic model of inbreeding depression that allows purging, is further justified because previous genetically-explicit modelling (Duthie & Reid, 2016) showed that inbreeding avoidance in biparental populations has a negligible effect on load given small-effect deleterious mutations (see also Wang *et al.*, 1999; Guillaume & Perrin, 2006).

We model inbreeding depression as having an absolute rather than relative effect on offspring viability (i.e., hard rather than soft selection) so that the effect of *β* is consistent across generations and different parameter combinations. We assume that inbreeding always decreases offspring viability (i.e., *β* > 0, giving inbreeding depression but no outbreeding depression). Therefore, because 0 ≤ *k*_*ij*_ ≤ 1, −*β* × *k*_*ij*_ *≤* 0. Values of Ψ_off_ must therefore be between zero (if −*β* × *k*_*ij*_ is very negative) and one (if −*β* × *k*_*ij*_ = 0). We therefore define Ψ_off_ as the probability that an offspring is viable, and sample its realised viability (versus mortality) using a Bernoulli trial. Offspring that are viable after the Bernoulli trial become adults. Given our current objectives, we simulate evolution under conditions where inbreeding avoidance is adaptive due to strong inbreeding depression, not where inbreeding preference is adaptive due to weak or zero inbreeding depression (Parker, 1979; Kokko & Ots, 2006; Duthie & Reid, 2016) or outbreeding depression (Bateson, 1983; Greeff *et al.*, 2009). However, as described above, positive *M*_*p*_ and *F*_*p*_ values resulting in inbreeding preference are not precluded from evolving, and could potentially arise due to mutation or drift.

### Immigration

After offspring mortality, *ρ* adult immigrants are added to the focal population. The kinship between an immigrant and all other individuals always equals zero (*k*_*ij*_ = 0). Immigration therefore prevents the mean kinship within the population from asymptoting to one over generations. To ensure that immigrants do not directly affect genotypic or phenotypic values of tendency for polyandry or pre-copulatory or post-copulatory inbreeding avoidance, immigrants are always male. Consequently, they can be chosen as females’ mates based on their values of *k*_*ij*_ = 0 but do not actively affect reproductive decisions through the expression of *P*_*p*_, *M*_*p*_, or *F*_*p*_. Further, immigrants’ *P*_*a*_, *M*_*a*_, and *F*_*a*_ allele values are randomly sampled from normal distributions with means and standard deviations equal to those in the focal population at the time of immigration, meaning that they do not directly cause any change in the distribution of allele values. We thereby effectively assume that the focal population receives immigrants from other nearby populations that are subject to the same selection on *P*_*p*_, *M*_*p*_, and *F*_*p*_ (Duthie *et al.*, 2016; Duthie & Reid, 2016).

### Density regulation

To avoid unrestricted population growth, we set separate carrying capacities for the total numbers of females (*K*_*f*_) and males (*K*_*m*_) in the focal population following immigration (Guillaume & Perrin, 2009; Duthie *et al.*, 2016). Hence, if at the end of a generation the number of females or males exceeds *K*_*f*_ or *K*_*m*_ respectively, then individuals are randomly removed until each sex is at its carrying capacity. Such removal can be interpreted as some combination of dispersal and mortality. The remaining females and males form the next generation of potentially breeding adults.

### Simulation and analysis

To address whether or not evolution of post-copulatory inbreeding avoidance alongside pre-copulatory inbreeding avoidance can facilitate evolution of polyandry, we compare simulations in which polyandry and pre-copulatory and post-copulatory inbreeding avoidance can all evolve with otherwise identical simulations in which post-copulatory inbreeding avoidance cannot evolve. To achieve this, we sever the connection from *F*_*a*_ to *F*_*p*_ such that all *F*_*g*_ genotypes cause random fertilisation with respect to kinship, so *F*_*a*_ alleles have no phenotypic effect. Simulations were repeated across four different costs of polyandry (*c*_*P*_ = {0, 0.0025, 0.005, 0.01}).

To address how asymmetric costs associated with pre-copulatory and post-copulatory inbreeding avoidance and polyandry affect the long-term persistence of reproductive strategies, we quantify the change in *M*_*a*_ and *F*_*a*_ over generations in simulations where pre-copulatory inbreeding strategy was cost free (*c*_*M*_ = 0) but post-copulatory inbreeding strategy was moderately costly (*c*_*F*_ = 0.02), and vice versa. We compare evolutionary trajectories with those of a costly strategy in the absence of evolution of an alternative strategy (e.g., evolution of pre-copulatory inbreeding strategy when post-copulatory inbreeding strategy phenotype is fixed at zero, *F*_*p*_ = 0). Previous modelling using similar genetic architecture suggests that a cost of 0.02 imposes strong but not overwhelming direct negative selection on polyandry (Duthie *et al.*, 2016). This value is therefore appropriate to illustrate the different evolutionary consequences that could result from asymmetrical costs. Results from simulations with additional cost value combinations are provided in Supporting Information.

To address how selection on an initially rare inbreeding avoidance strategy, and resulting evolution, is affected by the other strategy of inbreeding avoidance already being fixed in the population, we first used exploratory simulations to quantify evolution of pre-copulatory inbreeding strategy, and of post-copulatory inbreeding strategy and associated polyandry, in isolation. Then, to test whether pre-copulatory inbreeding avoidance would evolve when adaptive polyandry and post-copulatory inbreeding avoidance were fixed, we initiated *M*_*a*_ allele values at zero, but fixed *F*_*a*_ allele values at −10 and *P*_*a*_ allele values at 1 (i.e., *F*_*p*_ and *P*_*p*_ were expressed but did not evolve further). Similarly, to test whether post-copulatory inbreeding avoidance would evolve when pre-copulatory inbreeding avoidance was already fixed, we initiated *F*_*a*_ and *P*_*a*_ allele values at zero but fixed *M*_*a*_ allele values at −10. Consequently, because females have 10 diploid loci, when *M*_*a*_ or *F*_*a*_ alleles were fixed at −10, outbred females were 51 times less likely to choose a full brother and 13.5 times times less likely to choose a first cousin than a non-relative in pre-copulatory and post-copulatory choice, respectively.

In all simulations, we recorded mean values of *P*_*a*_, *M*_*a*_, and *F*_*a*_ in each generation and present these values over generations to infer selection on phenotypes (*P*_*p*_, *M*_*p*_, and *F*_*p*_). Each combination of parameter values simulated was replicated 40 times, and grand mean values and standard errors of means are calculated in each generation across replicates. These analyses allowed us to infer how allele values changed over generations in response to costs, but also in response to the changing values of other alleles and therefore potential evolutionary feedbacks among reproductive strategies. We do not use statistical tests to interpret simulation results; such tests are inappropriate because simulations violate key assumptions of statistical hypothesis testing, and statistical power (and therefore p-values) is determined entirely by the number of simulation replicates (White *et al.*, 2014).

For all replicates, we set the maximum number of generations to 40000, which exploratory simulations and previous modelling (Duthie *et al.*, 2016) showed to be sufficient for inferring long-term dynamics of mean allele values and therefore selection on phenotypes. For all replicates, we set *ρ* = 5 immigrants, which produced a range of kin and non-kin in each generation allowing females to express inbreeding strategies, and *n* = 8 offspring, which was sufficient to keep populations consistently at carrying capacities and avoid population extinction. Values of *K*_*f*_ and *K*_*m*_ were both set to 100 because previous modelling showed that populations of this size are small enough that mate encounters between kin occur with sufficient frequency for selection on inbreeding strategy, but not so small that selection is typically overwhelmed by drift (Duthie & Reid, 2016).

## Results

### Does evolution of post-copulatory inbreeding avoidance facilitate evolution of costly polyandry?

When post-copulatory inbreeding avoidance alleles (*F*_*a*_) had no effect (i.e., *F*_*g*_ values were fixed to zero), meaning that *F*_*p*_ could not evolve, *P*_*a*_ alleles underlying polyandry decreased to negative values over generations (red lines, Figure 1A,C,E,G). This shows that despite strong inbreeding depression in offspring viability, there is selection against unconditional polyandry even given zero direct cost (*c*_*P*_ = 0; Figure 1A). This is because *M*_*a*_ values became negative over generations, meaning that females typically avoided inbreeding through their initial mating (blue lines, Figure 1A,C,E,G). Polyandrous females that subsequently sampled more males from the available population were consequently more likely to mate with some relatives and hence produce some inbred offspring with low viability (see also Duthie *et al.*, 2016).

**Figure 1:**
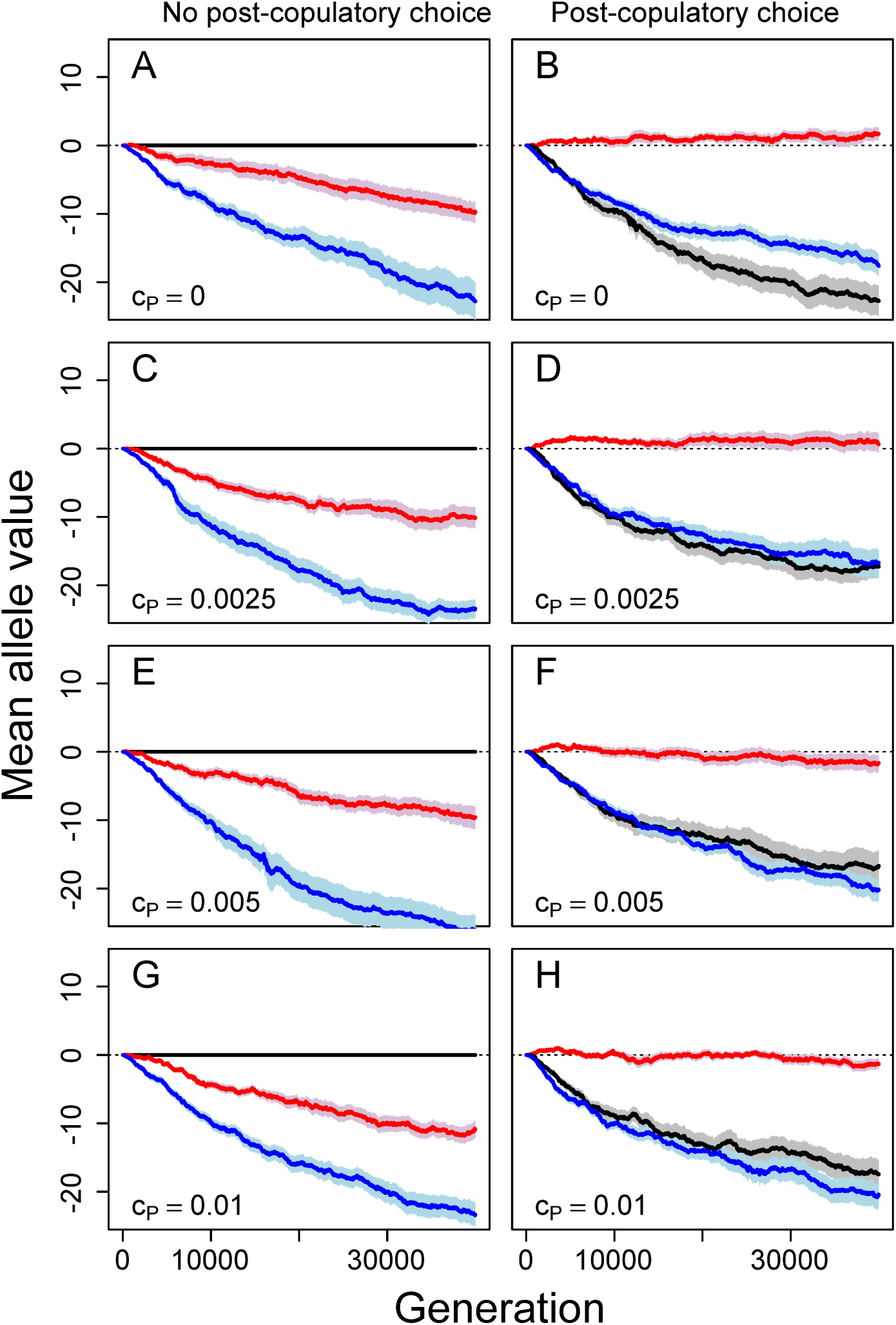
Mean allele values underlying tendency for polyandry (red), pre-copulatory inbreeding strategy (blue), and post-copulatory inbreeding strategy (black) from simulations where post-copulatory inbreeding strategy is (A, C, E, and G) fixed to zero (i.e., random fertilisation) or (B, D, F, and H) allowed to evolve freely. Costs of polyandry (*c*_*p*_) increase across rows from zero (A and B) to 0.01 (G and H). Mean allele values (solid lines) and associated standard errors (shading) are calculated across all individuals within a population over 40000 generations across 40 replicate populations. Negative mean allele values indicate inbreeding avoidance or tendency for monandry, and positive values indicate inbreeding preference or tendency for polyandry. Dotted lines demarcate mean allele values of zero.

When post-copulatory inbreeding avoidance was allowed to evolve, mean *P*_*a*_ values became substantially higher than in comparable simulations where *F*_*a*_ values were fixed to zero and post-copulatory inbreeding avoidance could not evolve (Figure 1B,D,F,H). Allowing evolution of post-copulatory inbreeding avoidance alongside pre-copulatory inbreeding avoidance therefore facilitated evolution of polyandry to the degree that most females mated multiply given low costs of polyandry (*c*_*P*_ < 0.005; e.g., Figure 2A,B). Here, *P*_*a*_ allele values increased from zero and persisted at low positive values (Figure 1B,D). Given higher costs of polyandry (*c*_*P*_ ≥ 0.005), *P*_*a*_ allele values still initially increased from zero, but then became slightly negative over generations (Figure 1F,H). Trajectories of allele values in individual simulations were typically highly stochastic, but were consistent in their long-term direction (Supporting Information Figures S1-S8). Overall, these results illustrate that post-copulatory inbreeding avoidance can facilitate evolution of polyandry as long as direct costs are sufficiently low (Figure 1B,D). However, given higher costs, evolution of polyandry is constrained even given strong inbreeding depression in offspring viability, and given resulting evolution of both pre-copulatory and post-copulatory inbreeding avoidance (Figure 1F,H).

**Figure 2:**
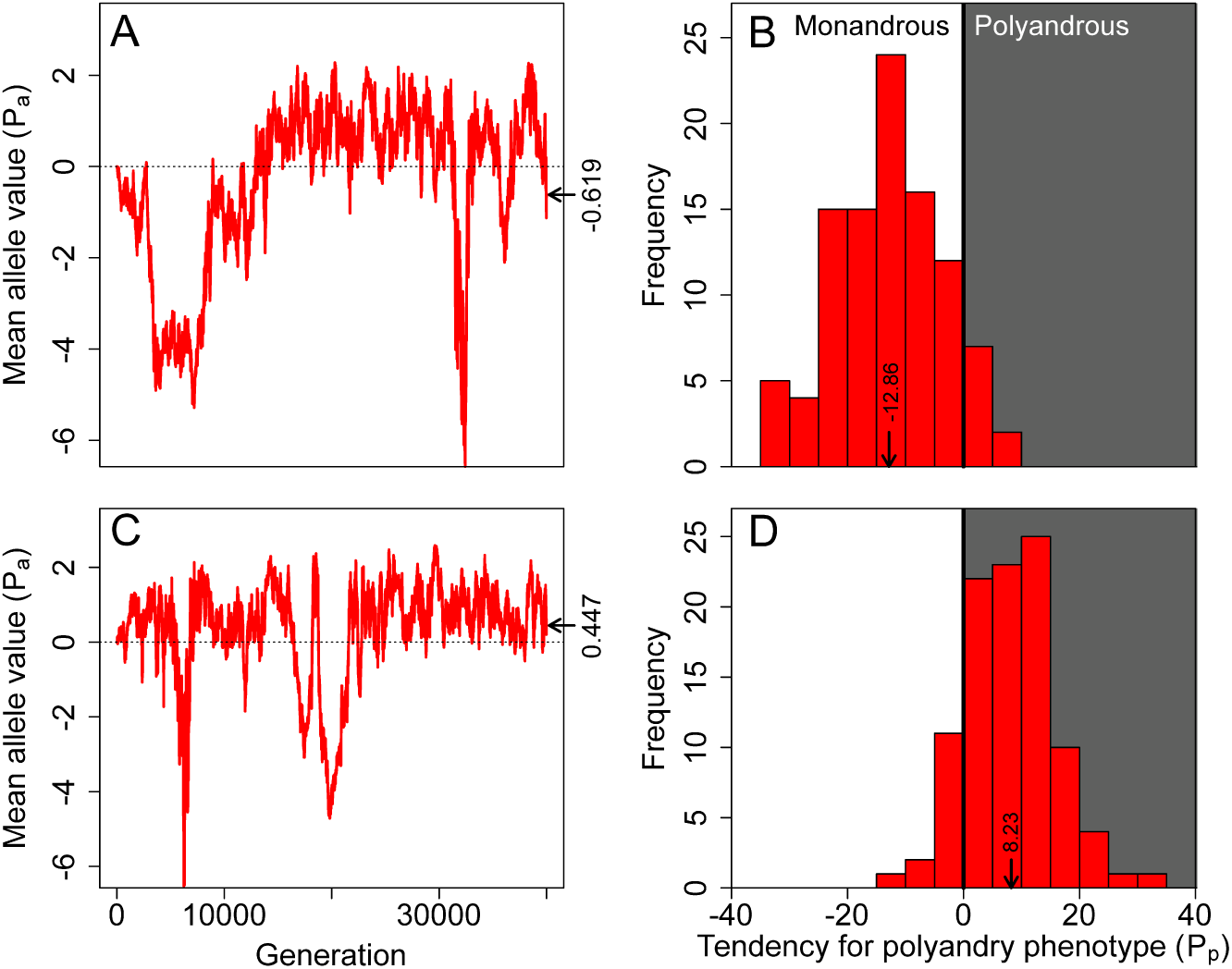
Relationships between (A and C) polyandry allele values and (B and D) monandry and polyandry phenotypes for simulations with identical initial conditions, default parameter values, and zero costs. Red lines in (A) and (C) show mean polyandry allele values across all individuals in a single simulation over 40000 generations. Positive and negative allele values contribute to polyandry and monandry, respectively. In the final generation, mean allele value was below (A) or above (C) zero (demarcated by the dotted line). Nevertheless, due to the threshold nature of expression of the polygenic polyandry phenotype, polyandry and monandry are expressed in both populations. Histograms in (B) and (D) show females’ tendency for polyandry phenotypes in the final generation; white and grey shading indicates monandrous and polyandrous females, respectively. Arrows and numbers indicate mean phenotype values. Because each trait includes 10 diploid loci with additive effects, phenotype values are ca 20 times allele values.

Strong post-copulatory inbreeding avoidance, manifested as very negative *F*_*a*_ values, consistently evolved in all simulations where such evolution was allowed (black lines, Figure 1B,D,F,H). Evolution of post-copulatory inbreeding avoidance occurred even when *P*_*a*_ values were expected to be slightly negative, and hence when there was selection against alleles underlying polyandry (Figure 1F,H). This reflects the threshold nature of phenotypic expression of polygenic polyandry, wherein random sampling of alleles means that polyandry is expressed by some females (i.e., *P*_*g*_ > 0) even when mean *P*_*a*_ values are negative (Figure 2). This means that, even in populations where female reproductive strategy evolves toward monandry, there is still commonly some opportunity for expression of post-copulatory inbreeding avoidance and associated selection that drives evolution of post-copulatory inbreeding avoidance.

Strong pre-copulatory inbreeding avoidance evolved (i.e., *M*_*a*_ < 0) in all simulations, irrespective of *c*_*P*_ and irrespective of whether post-copulatory inbreeding avoidance was allowed to evolve or hence whether polyandry evolved (Figure 1). This might be expected given strong inbreeding depression in offspring viability, which imposes selection against inbreeding.

### How do cost asymmetries affect long-term persistence of reproductive strategies?

When post-copulatory inbreeding avoidance allele values (*F*_*a*_) were fixed to zero, pre-copulatory inbreeding avoidance evolved even when costly (Figure 3A). Likewise, when pre-copulatory inbreeding avoidance allele values (*M*_*a*_) were fixed to zero, costly post-copulatory inbreeding avoidance evolved (Figure 3C). Females therefore evolved to avoid inbreeding, and thereby avoid the indirect cost of producing inbred offspring, through whichever route was available.

**Figure 3:**
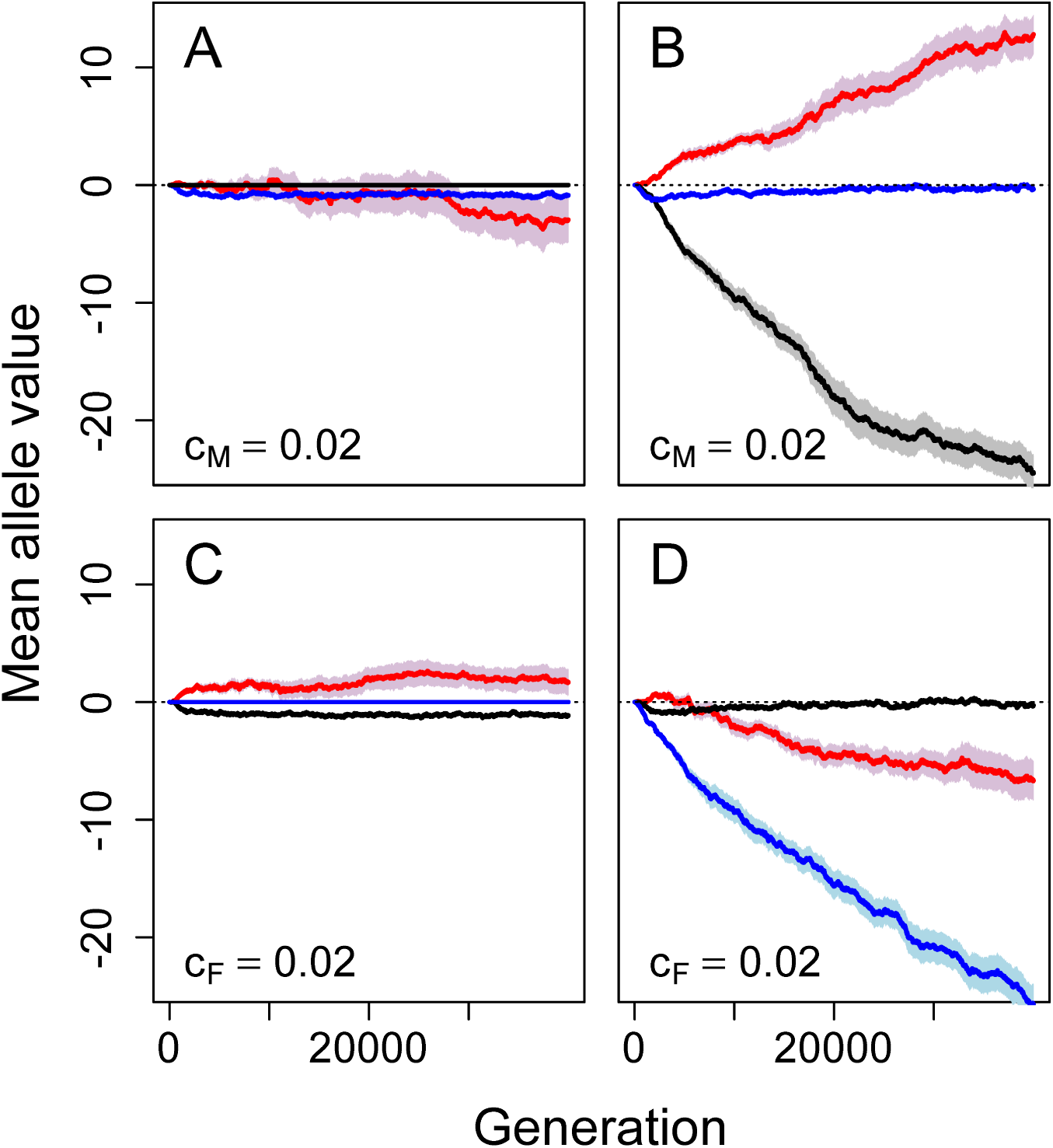
Mean allele values underlying tendency for polyandry (red), pre-copulatory inbreeding strategy (blue), and post-copulatory inbreeding strategy (black) when (A and B) costly pre-copulatory inbreeding strategy (*c*_*M*_ = 0.02) can evolve and post-copulatory inbreeding strategy is (A) fixed for random fertilisation or (B) can also evolve, and when (C and D) costly post-copulatory inbreeding strategy (*c*_*F*_ = 0.02) can evolve and pre-copulatory inbreeding strategy is (C) fixed for random mating or (D) can also evolve. Mean allele values (solid lines) and associated standard errors (shading) are calculated across all individuals within a population over 40000 generations across 40 replicate populations. Negative mean allele values indicate strategies of inbreeding avoidance or tendency for monandry, and positive values indicate strategies of inbreeding preference or tendency for polyandry. In all panels, polyandry is cost free.

However, when both pre-copulatory and post-copulatory inbreeding avoidance could evolve, their relative evolutionary dynamics depended on their relative costs. When pre-copulatory but not post-copulatory inbreeding avoidance was costly (*c*_*M*_ = 0.02 and *c*_*F*_ = 0), pre-copulatory inbreeding avoidance initially evolved (i.e., *M*_*a*_ < 0) but then evolved back towards random mating (i.e., *M*_*a*_ ≈ 0) following increasing evolution of post-copulatory inbreeding avoidance and polyandry (Figure 3B). Similarly, when post-copulatory but not pre-copulatory inbreeding avoidance was costly (*c*_*F*_ = 0.02 and *c*_*M*_ = 0), post-copulatory inbreeding avoidance initially evolved (i.e., *F*_*a*_ < 0) before evolving back to random fertilisation (i.e., *F*_*a*_ ≈ 0) after ca 20000 generations (Figure 3D).

Further simulations illustrate that such evolution of a high cost inbreeding strategy back toward random mating or random fertilisation in the presence of a relatively low cost alternative strategy is a general consequence of differential selection on each strategy induced by cost asymmetry within a range of relatively small costs, not specific to values of *c*_*F*_ and *c*_*M*_ of 0 and 0.02. For example, cost-specific evolution of pre-copulatory or post-copulatory inbreeding avoidance also occurred given non-zero small costs (e.g., *c*_*F*_ and *c*_*M*_ values of 0.01 and 0.03 and vice versa; Figure S9) and smaller cost asymmetries (e.g., *c*_*F*_ and *c*_*M*_ values of 0.01 and 0.02 and vice versa; Figure S10).

When pre-copulatory inbreeding avoidance was costly, allowing evolution of cost free post-copulatory inbreeding avoidance greatly facilitated evolution of polyandry (Figure 3A versus 3B). However when post-copulatory inbreeding avoidance was costly, allowing evolution of cost free pre-copulatory inbreeding avoidance caused *P*_*a*_ alleles to decrease to very negative values, reducing expression of polyandry (Figure 3C versus 3D; polyandry was cost free in all these simulations). Results for all possible cost combinations of 0 and 0.02, including costly polyandry, are provided in Supporting Information (Figure S11).

### How does fixation of pre-copulatory or post-copulatory inbreeding avoidance affect evolution of an alternative strategy of inbreeding avoidance?

When polyandry alleles (*P*_*a*_) were fixed to be positive so that all females were expected to mate multiply and post-copulatory inbreeding allele (*F*_*a*_) values were fixed for adaptive inbreeding avoidance, pre-copulatory inbreeding avoidance evolved (i.e., *M*_*a*_ values became increasingly negative; Figure 4A). Such evolution still occurred, but to a much smaller degree, when pre-copulatory inbreeding avoidance was costly (Figure 4B). However, after 40000 generations, *M*_*a*_ values were less negative when post-copulatory inbreeding avoidance and polyandry were fixed at non-zero values than when they also evolved from initial values of zero (–15.45 vs. –17.62; compare the blue lines in Figures 4A versus Figure 1B). This implies that pre-existence of fixed post-copulatory inbreeding avoidance can weaken selection and subsequent evolution of pre-copulatory inbreeding avoidance, but does not necessarily preclude it.

**Figure 4:**
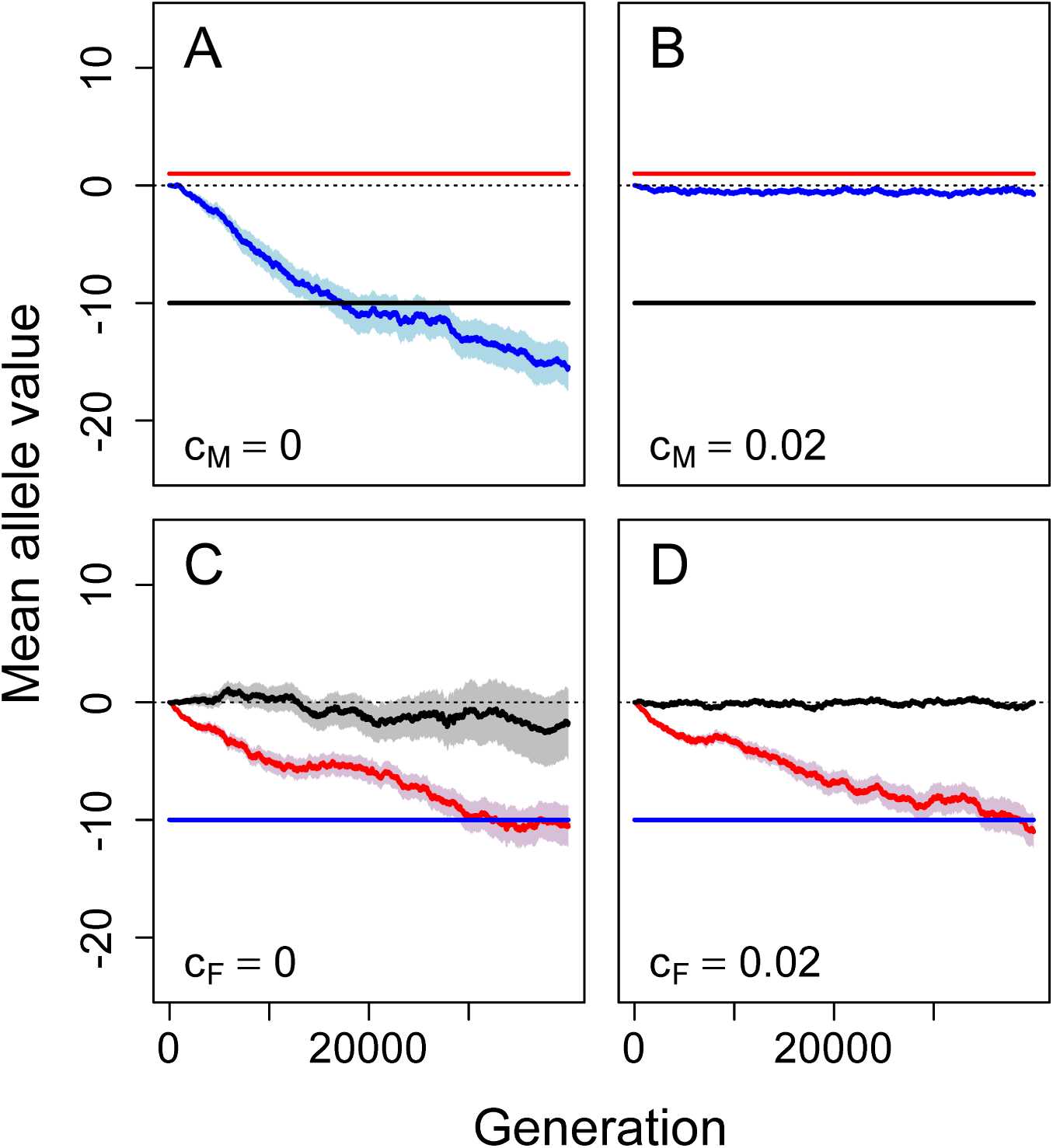
Mean allele values underlying tendency for polyandry (red), pre-copulatory inbreeding strategy (blue), and post-copulatory inbreeding strategy (black), given (A and B) fixed polyandry and post-copulatory inbreeding avoidance and (C and D) fixed post-copulatory inbreeding avoidance where the evolving inbreeding strategy is cost-free (A, *c*_*M*_ = 0; C, *c*_*F*_ = 0) or costly (B, *c*_*M*_ = 0.02; D, *c*_*F*_ = 0.02). Mean allele values (solid lines) and associated standard errors (shading) are calculated across all individuals within a population over 40000 generations across 40 replicate populations. Negative mean allele values indicate strategies of inbreeding avoidance or tendency for monandry, and positive values indicate strategies of inbreeding preference or tendency for polyandry.

In contrast, when pre-copulatory inbreeding allele (*M*_*a*_) values were fixed for adaptive inbreeding avoidance, mean *F*_*a*_ allele values did not consistently become negative over generations (Figure 4C,D). This implies that existence of fixed pre-copulatory inbreeding avoidance can prevent evolution of post-copulatory inbreeding avoidance. In these simulations, mean *P*_*a*_ allele values consistently decreased over generations, reflecting selection against polyandry regardless of whether or not post-copulatory inbreeding avoidance was costly (Figure 4C,D). Consequently, when *c*_*F*_ = 0, *F*_*a*_ allele values had no effect because females were almost exclusively monandrous, resulting in high drift of *F*_*a*_ values (resulting in variation among replicates illustrated by the wide standard errors in Figure 4C). However, when *c*_*F*_ = 0.02, *F*_*a*_ values remained near zero to minimise direct costs. The lack of selection for post-copulatory inbreeding avoidance when pre-copulatory inbreeding avoidance was fixed was driven by a lack of polyandry, and therefore an inability of females to bias fertilisation among multiple mates. When pre-copulatory inbreeding avoidance and polyandry were both fixed (*M*_*a*_ = –10 and *P*_*a*_ = 1), *F*_*a*_ allele values evolved to similarly negative means as *M*_*a*_ allele values in Figure 4A,B (see Supporting Information Figure S12).

## Discussion

Different reproductive strategies cannot be presumed to evolve in isolation from one another. Rather, there is likely to be considerable potential for feedbacks and degeneracy (i.e., functional redundancy) among interacting phenotypes. For example, inbreeding avoidance could be manifested through both pre-copulatory and post-copulatory mechanisms and associated polyandry, meaning that simultaneous evolution of each phenotype might be affected by degeneracy, in addition to trait-specific benefits and costs.

We used individual-based modelling to highlight fundamental but theoretically under-developed relationships between evolution of polyandry and pre-copulatory versus post-copulatory inbreeding strategy given (1) hard constraints on evolution of post-copulatory inbreeding strategy, (2) asymmetric costs of pre-copulatory and post-copulatory inbreeding strategy, and (3) evolution of one inbreeding strategy phenotype given pre-existence of the other. Our current model and simulation results thereby provide tools for thinking more clearly about the dynamics of simultaneously evolving reproductive strategies in the context of polyandry and inbreeding avoidance.

### Interacting evolution of polyandry and inbreeding avoidance strategies

The opportunity to adjust inbreeding is widely suggested to be a driver of adaptive evolution of polyandry (Tregenza & Wedell, 2002; Foerster *et al.*, 2003; Akçay & Roughgarden, 2007; Varian-Ramos & Webster, 2012; Kingma *et al.*, 2013; Lehtonen & Kokko, 2015; Reid *et al.*, 2015*a*). Our simulations show that when post-copulatory inbreeding avoidance could evolve, selection for and resulting evolution of polyandry was greatly strengthened (Figure 1). The proposition that polyandry might facilitate cryptic female choice among males of varying compatibility is not new (e.g., Zeh & Zeh, 1997; Jennions & Petrie, 2000), but our model clarifies this verbal hypothesis and therefore has widespread implications for future studies of polyandry evolution.

We predict evolution of polyandry in populations where inbreeding depression is severe and inbreeding avoidance through post-copulatory mechanisms can also evolve, especially if pre-copulatory inbreeding avoidance is costly (Figure 1). Indeed, post-copulatory inbreeding avoidance has been observed under these conditions in experimental systems across diverse taxa (e.g., Pizzari *et al.*, 2004; Firman & Simmons, 2008, 2015; Bretman *et al.*, 2009; Gasparini & Pilastro, 2011; Tuni *et al.*, 2013).

Evolution of both pre-copulatory and post-copulatory inbreeding avoidance occurred in our model, but were affected by the evolution of polyandry and by cost asymmetries. One cost free strategy of inbreeding avoidance precluded another more costly strategy from persisting in a focal population (Figure 3). Hence, our model demonstrates degeneracy between inbreeding avoidance strategies, and implies that such interactions should be considered when developing hypotheses concerning reproductive strategy within and across systems. In particular, indefinite persistence of both pre-copulatory and post-copulatory inbreeding avoidance should not be expected in populations given a sufficiently large and sustained cost asymmetry. However, the time required for the more costly inbreeding strategy to go extinct might be on the order of tens of thousands of generations (Figure 3), and spatial or temporal variation in costs might facilitate coexistence of multiple inbreeding avoidance strategies.

Further, pre-existence of fixed adaptive pre-copulatory inbreeding avoidance precluded evolution of polyandry and, in turn, precluded evolution of post-copulatory inbreeding avoidance (Figure 3C,D). However, pre-existence of fixed adaptive post-copulatory inbreeding avoidance did not preclude evolution of pre-copulatory inbreeding avoidance (Figure 3A,B). In natural populations, it is unlikely that pre-copulatory and post-copulatory inbreeding avoidance will evolve simultaneously from an ancestral population in which females mate and assign paternity randomly. Rather, the timing of the invasion of adaptive inbreeding avoidance phenotypes will differ, so the initial evolution of one inbreeding strategy will likely occur in the absence of the other, or where selection for and subsequent evolution of the other strategy has already occurred. When framing hypotheses for existence of post-copulatory inbreeding avoidance and polyandry, it might therefore be necessary to consider whether or not inbreeding avoidance is already known to occur through pre-copulatory mate choice. Additionally, the opportunity for post-copulatory inbreeding avoidance will also depend on the degree to which females are polyandrous. For species in which pre-copulatory inbreeding avoidance occurs and polyandry is uncommon (Lihoreau *et al.*, 2007; Metzger *et al.*, 2010*a*,*b*), evolution of post-copulatory inbreeding avoidance is unlikely even if such a strategy incurs little direct cost.

### General hypotheses concerning inbreeding avoidance and polyandry

Post-copulatory inbreeding avoidance cannot be effectively realised if females are not polyandrous in any form, and is likely to be most effective for highly polyandrous females that can bias fertilisation amongst sperm contributed by multiple mates. In contrast, pre-copulatory inbreeding avoidance mechanisms are most critical for females that mate only once and therefore have no post-copulatory opportunity to avoid inbreeding. This theory is borne out in our simulation results, as selection for, and consequent evolution of, post-copulatory inbreeding avoidance was negligible in populations where polyandry did not evolve, resulting in high drift of allele values over generations due to the inability of females to express post-copulatory choice (e.g., Figure 4C). Evolution of pre-copulatory inbreeding avoidance was also typically slightly stronger when polyandry did not evolve (e.g., Figure 3A vs 3B; see also Supporting Information Figure S11). In addition to initial polyandry causing evolution of post-copulatory inbreeding avoidance, polyandry might also covary positively with post-copulatory inbreeding avoidance due to the feedback effect that post-copulatory inbreeding avoidance has on facilitating evolution of polyandry itself, as observed in our model (Figure 1). It would therefore be interesting to test the hypothesis that, across taxa, the occurrence of post-copulatory inbreeding avoidance covaries positively, and the occurrence of pre-copulatory inbreeding avoidance covaries negatively, with the degree of polyandry. To test this hypothesis, further empirical work is needed to quantify the degree to which females of different species engage in polyandry and the degree to which females express both pre-copulatory and post-copulatory inbreeding avoidance.

Degeneracy occurs at nearly all biological scales (Edelman & Gally, 2001), including complex systems affecting organismal development (e.g., Nowak *et al.*, 1997), adaptation (Whitacre & Bender, 2010; Whitacre, 2010), and cognition (Price & Friston, 2002; Park & Friston, 2013), as well as population (Atamas & Bell, 2009), community (Suraci *et al.*, 2017), and ecosystem (e.g., Levin & Lubchenco, 2008) dynamics. In our model, degeneracy occurred through overlaps in how different reproductive strategies caused adaptive inbreeding avoidance. In general, degeneracy might increase biological robustness by fine-tuning degenerate phenotypes to different local environments (Gardner & Kalinka, 2006; Whitacre, 2010). For example, degeneracy might ensure successful inbreeding avoidance through either pre-copulatory or post-copulatory mechanisms when avoidance through the other mechanism is ineffective (e.g., due to sexual conflict affecting mate choice or injury affecting fertilisation). However, evolution of one inbreeding avoidance mechanism might also weaken selection on the other by modifying the latter’s impact on total realised inbreeding avoidance (*sensu* evolution of genetic redundancy; see Nowak *et al.*, 1997). The relevance of degeneracy with respect to such reproductive strategies therefore requires further theoretical development, which could result in new empirical predictions and conceptual synthesis across biological scales.

### Model structure, assumptions, and extensions

While the logic of our current model can be usefully applied to construct general hypotheses within and across empirical systems, accurate quantitative prediction for specific systems would require additional empirical detail and data for model parameterisation. To facilitate general conceptual comparison of the effects of direct costs across phenotypes, we modelled all costs as analogous increased probabilities of female mortality and hence total reproductive failure. This cost formulation reflects empirical observations in some populations (see Model; e.g., Rowe *et al.*, 1994; Koga *et al.*, 1998; Gasparini & Pilastro, 2011), and is therefore a biologically realistic method of standardising costs across traits. However, different forms of costs could be incorporated into future models designed to predict specific evolutionary dynamics. Models could then be further developed such that costs arise from explicit reproductive mechanisms, requiring further biological detail.

For example, Pomiankowski (1987) identified four cost categories relevant to mating frequency and mate choice, including elevated risks of predation or disease transmission, and time or energy expenditure. Polyandrous females might experience increased risk of disease transmission (Roberts *et al.*, 2015), a cost that would more realistically apply to a female’s realised number of mates rather than her tendency for polyandry. Polyandrous females might also risk harm caused by sexual conflict over multiple mating (e.g., Arnqvist & Rowe, 2005; Parker, 2006). Inbreeding theory predicts that males should be more tolerant of inbreeding than females, leading to sexual conflict over inbreeding in mating encounters (Parker, 1979, 2006; Kokko & Ots, 2006; Duthie & Reid, 2015). Future models could therefore explicitly consider sexual conflict over both polyandry and pre-copulatory inbreeding, and hence capture internally-consistent mechanistic costs. Further, sexual conflict might also affect post-copulatory inbreeding avoidance. For example, when female guppies (*Poecilia reticulata*) were artificially inseminated with equal quantitites of sperm from full-siblings and unrelated males, more eggs were fertilised by unrelated males because the velocities of full sibling’s sperm were reduced by females’ ovarian fluids (Gasparini & Pilastro, 2011). In black field crickets (*Teleogryllus commodus*), females attempt to remove the spermatophores of unwanted males after copulation, and are capable of controlling sperm transfer to spermatheca after copulation occurs (Bussiére *et al.*, 2006; Tuni *et al.*, 2013). Future models could therefore explicitly incorporate such mechanisms in order to better understand effects of post-copulatory sexual conflict on female and male reproductive strategy evolution.

We restricted our current model to examine how reproductive strategy evolution was affected by direct costs and indirect benefits stemming from inbreeding avoidance, thereby isolating such effects and explicitly addressing key general hypotheses regarding the effects of inbreeding depression. In natural populations, reproductive strategy evolution will also be affected by other costs and benefits, for example, stemming from additive genetic effects (i.e., pre-copulatory or post-copulatory mate choice for ‘good genes’). Future objectives could consequently be to develop theory and models that include multiple benefits and costs of reproductive strategies, which could then be parameterised using empirical data. While good theory should always strive for conceptual clarity and the avoidance of unnecessary nuance, the prudent use of multi-faceted mechanistic models could usefully link theory, modelling, and empirical hypothesis testing and thereby improve both general and specific understanding and prediction of reproductive strategy evolution.

## Acknowledgments

This work was funded by a European Research Council Starting Grant to JMR. Computer simulations were performed using the Maxwell Computing Cluster at the University of Aberdeen. We thank Matthew E. Wolak for very helpful comments.

